# Assessing horizontal gene transfer in the rhizosphere of *Brachypodium distachyon* using fabricated ecosystems (EcoFABs)

**DOI:** 10.1101/2024.03.14.584828

**Authors:** Shweta Priya, Silvia Rossbach, Thomas Eng, Hsiao-Han Lin, Peter F. Andeer, Jenny C. Mortimer, Trent R. Northen, Aindrila Mukhopadhyay

**Affiliations:** Biological Systems and Engineering Division, Lawrence Berkeley National Laboratory, Berkeley, CA, 94720, USA; Environmental Genomics and Systems Biology Division, Lawrence Berkeley National Laboratory, Berkeley, CA, 94720, USA; Department of Biological Sciences, Western Michigan University, 1903 W Michigan Ave, Kalamazoo MI 49008-5410, U.S.A; School of Agriculture, Food and Wine, University of Adelaide, Australia

## Abstract

Horizontal gene transfer (HGT) is a major process by which genes are transferred within microbes in the rhizosphere. However, examining HGT remains challenging due to the complexity of mimicking conditions within the rhizosphere. Fabricated ecosystems (EcoFABs) have been used to investigate several complex processes in plant associated environments. Here we show that EcoFABs are efficient tools to examine and measure HGT frequency in the rhizosphere. We provided the first demonstration of gene transfer via triparental conjugation system in the *Brachypodium distachyon* rhizosphere in the EcoFABs using *Pseudomonas putida* KT2440 as both donor and recipient bacterial strain with the donor having the mobilizable and non-self-transmissible plasmid. We also observed that the frequency of conjugal plasmid transfer in the rhizosphere is potentially dependent on the plant developmental stage, and composition and amount of root exudates. The frequency of conjugation also increased with higher numbers of donor cells. We have also shown the transfer of plasmid from *P. putida* to another *B. distachyon* root colonizer, *Burkholderia* sp. showing the possibility of HGT within a rhizosphere microbial community. Environmental stresses also influence the rate of HGT in the rhizosphere between species and genera. Additionally, we observed transfer of a non-self transmissible donor plasmid without the helper strain on agar plates when supplemented with environmental stressors, indicating reduced dependency on the helper plasmid under certain conditions. This study provides a robust workflow to evaluate conjugal transfer of engineered plasmids in the rhizosphere when such plasmids are introduced in a field or plant associated environment.

**Importance:** We report the use of EcoFABs to investigate the HGT process in a rhizosphere environment. It highlights the potential of EcoFABs in recapitulating the dynamic rhizosphere conditions as well as their versatility in studying plant-microbial interactions. This study also emphasizes the importance of studying the parameters impacting the HGT frequency. Several factors such as plant developmental stages, nutrient conditions, number of donor cells and environmental stresses influence gene transfer within the rhizosphere microbial community. This study paves the way for future investigations into understanding the fate and movement of engineered plasmids in a field environment.

## 1. Introduction

Horizontal gene transfer (HGT) enables transmission of genetic material between otherwise unrelated organisms via transformation, transduction or conjugation. This process allows an organism to acquire new functions that can shape bacterial evolution or fitness by providing new genes that would be unlikely to arise from simpler spontaneous mutations. HGT can lead to the appearance of multiple identical, redundant operons within the soil communities which promote intra- and interspecies variability [1,2]. Conjugation is one of the major HGT processes by which bacteria acquire DNA and new functionality to adapt and overcome environmental stresses and compete in their ecological niches [3]. Non-conjugative plasmids from donor cells can be mobilized to recipient cells by helper plasmids; this process is known as triparental conjugation [4].

Such gene transfer phenomena are equally important to examine, but less understood in the context of genetically modified microbes and their potential use or release into the environment [2]. Specifically, HGT remains challenging to study and quantify in natural environments like the rhizosphere or in the soil matrix. Recently, lab-scale microcosms such as rhizotrons, rhizobox, nylon soil pouches etc., have been designed that allow continuous monitoring throughout the plant developmental stages and non-destructive sampling of the rhizosphere or root metabolites, in real-time [5]. Of these, fabricated ecosystems (EcoFABs) are open-source 3D printable chambers that are highly standardized reproducible tools enabling mechanistic studies of plant-microbe interactions under controlled laboratory conditions (**Fig. 1**) [6,7]. Several complex processes occurring in the rhizosphere under natural environmental conditions have been simulated and studied in EcoFABs [6,8,9]. In such studies, repeatability is critical. Sasse et al. (2019) found that *Brachypodium distachyon* growth in EcoFABs was reproducible across four laboratories for a number of morphological and metabolic traits of root tissue and root exudates [8]. EcoFABs may also enable measurement of gene transfer frequency under controlled conditions and mimic key aspects of the natural rhizosphere environment [7].

**Fig. 1.**
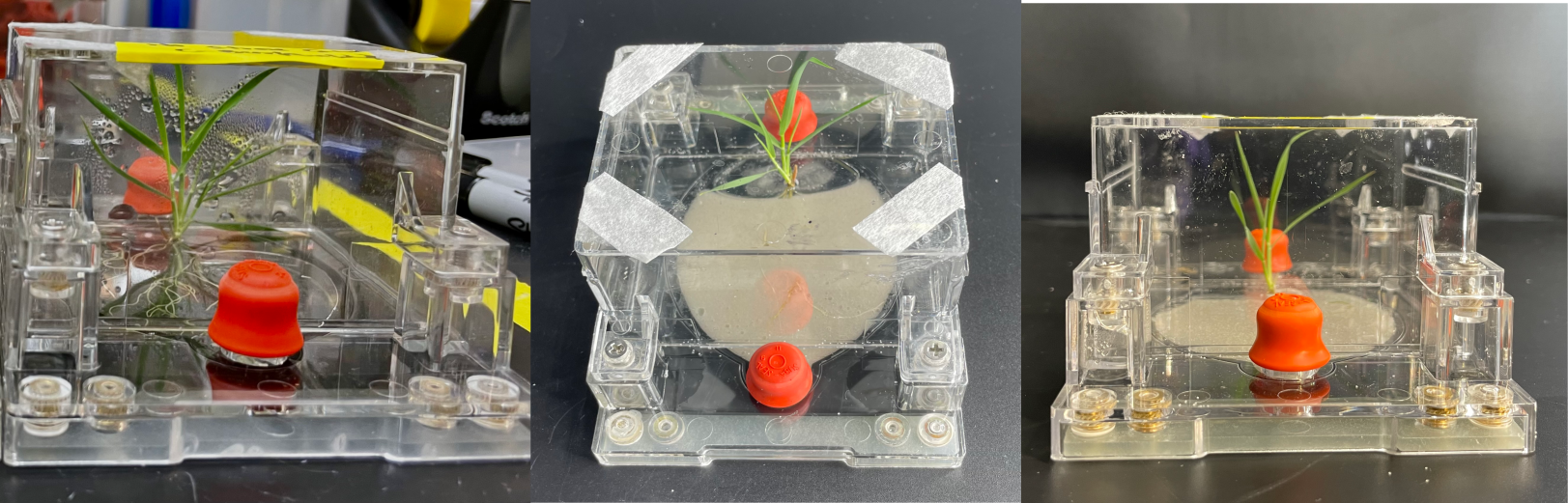
*Brachypodium distachyon* grown in EcoFAB 2.0 (microcosms used in this study). Pictures of EcoFAB taken with liquid medium, and sand from above and side respectively (left to right).

The frequency of gene transfer and uptake is potentially enhanced by availability of nutrients in the rhizosphere from root exudates, and other favorable conditions such as moisture and root surface that make the rhizosphere a preferred site for colonization [10]. The high cell densities and metabolic activity of microbes in the rhizosphere may also influence the HGT processes [11,12]. In addition to nutrient content, environmental stresses such as drought or salinity could also be the driving conditions for rhizospheric bacteria to acquire genes and plasmids from others in the community to compete and survive [13]. To our knowledge, there are few studies that have investigated the impact of abiotic stress conditions on the frequency of HGT but this could be a critical parameter influencing this phenomenon [14,15]. For instance, studies have reported that the rate of conjugal transfer of plasmids is influenced by changes in soil physico-chemical conditions such as moisture or osmotic stress, pH, temperature, and soil components which indicate that changes in rhizosphere conditions due to these abiotic stresses could also impact the HGT frequency [16,17]. While some studies have reported possible transmission of plasmids between bacteria of distant genera, this process is yet to be investigated in detail in the rhizosphere environment [16,18].

Here, we develop a model system with established HGT efficiencies to monitor plasmid uptake by root associated bacteria in the EcoFAB root chambers and measure HGT. We describe a triparental conjugation mating system inoculated onto *B. distachyon* roots. Establishing this baseline allowed assessment of other parameters such as donor to recipient ratio and environmental stresses that may impact the conjugation frequency in the rhizosphere. Furthermore, we also examined intergeneric plasmid transfer using a non-model bacterium of the rhizosphere microbial community as a recipient. To understand the possibility of transfer of a mobilizable and non-self transmissible plasmid without a helper strain, we have also tested bi-parental mating system under routine laboratory or environmentally stressed conditions.

## 2. Materials and Methods

### 2.1 Growing plants in EcoFABs

All experiments were performed with *B. distachyon* Bd21-3 as the host plant [19]. Seeds were dehusked and sterilized in 70% v/v ethanol for 30 s, and in 6% v/v sodium hypochlorite for 5 min, followed by five wash steps in sterilized water and kept at 4 °C under dark conditions for stratification for 7 days. Seedlings were germinated on 0.5x Murashige & Skoog (MS) plates (2.2 g l^−1^ MS medium, MSP01 (Caisson Labs, Smithfield, UT, USA) with 1650 mg l^−1^ ammonium nitrate, 6.2 mg l^−1^ boric acid, 332.2 mg l^−1^ calcium chloride, 0.025 mg l^−1^ cobalt chloride, 0.025 mg l^−1^ copper sulfate, 37.26 mg l^−1^ disodium EDTA, 27.8 mg l^−1^ ferrous sulfate heptahydrate, 180.7 mg l^−1^ magnesium sulfate, 16.9 mg l^−1^ manganese(II) sulfate monohydrate, 0.25 mg l^−1^ sodium molybdate dihydrate, 0.83 mg l^−1^ potassium iodide, 1900 mg l^−1^ potassium nitrate, 170 mg l^−1^ monopotassium phosphate, 8.6 mg l^−1^ zinc sulfate heptahydrate; 6% w/v phytoagar (Fisher Scientific, Waltham, MA, USA), pH adjusted to 5.8) in a 16 h : 8 h, light : dark regime at 24 °C with 150 µmol m^−2^ s^−1^ illumination in reach-in plant growth chambers (Controlled Environments Inc., Pembina, North Dakota, USA) [20]. EcoFAB 2.0 [6] were sterilized and seedlings transferred to EcoFAB chambers at 3 d after germination. The EcoFAB root chambers were filled with 10 g of sand (Sand, 50-70 mesh particle size, Sigma-Aldrich, St. Louis, MO) and 5 ml of 0.5 x MS before transferring to a humidity-controlled growth chamber. EcoFABs with plants were incubated in the plant growth chambers under the same conditions as mentioned above for seed germination. To impose osmotic stress conditions, 0.5x MS was added with 10% and 20% polyethylene glycol (PEG 6000, Sigma-Aldrich, St. Louis, MO) for low and high stress levels respectively before adding the media in the EcoFABs [21]. For salinity stress, 0.5 x MS was supplemented with 50 mM and 100 mM NaCl (Sigma-Aldrich, St. Louis, MO).

### 2.2 Harvesting and collection of exconjugants from rhizosphere

Plants were harvested at 10 days after seed emergence (DAG), 20 DAG and 30 DAG for the purpose of root exudate collection for HPLC (see section 2.3) and to examine the effect of developmental stage of plant on conjugation frequency. For all other experiments, plants were harvested after 2 weeks of growth (∼5 primary leaves under optimum conditions) and 24 hours post-inoculation. To examine if the presence of roots influence the conjugation frequency, we removed the roots from 3 EcoFABs (two weeks old plants) using sterile forceps after unscrewing the EcoFAB from top under aseptic conditions and dissolved the roots in 1mL sterile water, vortexed for 10 minutes to dissolve all the root exudates attached in the roots and this slurry was added back into the same EcoFABs and mixed well using a sterile spatula for uniform distribution of root exudates. All other EcoFABs for this set of experiments (results in section 3.1 and 3.2) were also watered with 1mL sterile water simultaneously.

All the EcoFABs (except those which were grown for root exudate collection for HPLC) were then inoculated and incubated for 24 hours (see section 2.5) before the harvesting day. The incubated EcoFABs were then opened under aseptic conditions to extract the conjugants. The EcoFABs containing inoculated *Brachypodium*, were unscrewed from the top and the roots were cut from the shoots using sterile blades and pulled out using sterile tweezers and the sand along with the roots (rhizosphere) was transferred to the 50 mL sterile centrifuge tubes containing 1 mL 1 x phosphate buffered saline (PBS). The tubes containing the roots in the PBS were shaken for 10 minutes to separate the attached microbes of the rhizosphere into the suspension. The suspension was then pipetted into a microcentrifuge tube, serially diluted and plated as detailed in next few sections.

### 2.3 Extraction of root exudates and quantification of sugars and organic acids

For root exudate collection, plants were grown separately for 10 DAG, 20 DAG and 30 DAG of seeds in replicates in the EcoFABs following the same procedure as section 2.1. EcoFABs filled with 10g sand (no plant) and 5 mL of 0.5 x MS were used as controls. The sand along with the roots (without roots for controls) were separated from shoots and transferred into 50 mL sterile centrifuge tubes containing sterile 10 mL of sterile ultrapure water and gently stirred for 30 minutes. The sand was settled down and the aliquot was filter-sterilized using a 0.22 μm filter, and lyophilized. The freeze dried root exudates were dissolved in 500 µ`L of sterile HPLC grade water and stored at −20 °C for downstream applications`. Amount of sugars and organic acids in the root exudates were quantified using Agilent HPLC 1260 infinity system (Santa Clara, CA, USA) equipped with a Bio-Rad Aminex HPX-87H column and a Refractive Index detector. An aqueous solution of sulfuric acid (4 mM) was used as the eluent (0.6 mL min^−1^, column temperature 60 °C).

### 2.4 Strains and plasmids used in the study

A uracil auxotroph *Pseudomonas putida* KT2440 Δ*pyrF* harboring the plasmid pSRLBL01 was used as the plasmid donor. This plasmid was used as a proxy for any commonly used engineered plasmid and carries the common genes that are the backbone of developing engineered microorganisms. The donor plasmid contains genes for gentamicin resistance as well as *cfp* encoding the cyan fluorescent protein. The recipient used was *P. putida* KT2440 with a chromosomally integrated mCherry gene. This experimental set-up allowed us to counterselect for the donor strain and to visualize any gene transfer based on fluorescence. To improve the conjugation frequency of the mobilizable, non self-transmissible donor plasmid, we included a helper strain *E. coli* DH5ɑ harboring the plasmid pRK2013 containing the *tra* genes that aided the transfer of donor plasmid to the recipient strain via a triparental conjugation system. This plasmid has been widely used as a conjugative helper plasmid to assist the transfer of a mobilizable and non-self transmissible plasmid from one bacteria to another [22,23]. For the intergeneric conjugation experiment, we used *Burkholderia sp.* OAS925 as recipient of the same donor plasmid.

**Table 1.** Plant, bacterial strains and plasmids used in the study.

| Plant or strain or plasmid | Genotype or phenotype | Reference |
| --- | --- | --- |
| *Brachypodium distachyon* BD21-3 | Plant | Vogel and Hill, 2008 [19] |
| *Pseudomonas putida* KT2440 Δ*pyrF* (JBEI-204817) | Donor strain, uracil auxotroph | Czajka et al., 2022 [24] |
| *P. putida* KT2440 PP_0871:pTE451 {Pglut-riboswitch-mCherry PclpB-mCitrine} KanS SucR (JBEI-255664) | Recipient strain | This study |
| *E.coli* DH5α (pRK2013) | Tra<sup>+</sup> Mob<sup>+</sup> (RK2) Km::Tn7 ColEI origin, helper plasmid, KmR | Figurski & Helsinki, 1979 [25] |
| *Burkholderia sp.* OAS925 | Recipient strain | Price et al., 2022 [26] |
| pSLRLBL01 (JBEI-255655) | Donor plasmid GntR, pBAD *cfp* | This study |

### 2.5 Preparation of bacterial cultures for triparental conjugation

The recipient strain was grown in LB broth and donor strain was grown in LB broth with 0.2 um filter sterilized 25 µg/mL Uracil (Sigma-Aldrich, St. Louis, MO) supplemented with 30 mg/L gentamicin and both strains were incubated overnight at 30 ℃ and 200 RPM. The helper strain was grown in LB medium containing 50 mg/L of kanamycin and incubated overnight at 37 ℃ and 200 RPM. The bacterial cells grown overnight were harvested by centrifuging at 4,000 *g* for 10 min and washed twice with sterile phosphate-buffered saline (PBS). The cell pellet was resuspended in sterile PBS and the OD_600_ was adjusted to obtain ∼10^5^ CFUs/mL of each strain.

### 2.6 Conjugation experiment in EcoFABs

To determine the conjugation frequency in the rhizosphere, *B. distachyon* plants grown in the EcoFABs (see section 2.1) were inoculated with the mixture of all three strains. Bacterial donor, recipient and helper cells were mixed in a microcentrifuge tube in the ratio of 10:1:1 respectively to a total volume of 100 uL. To evaluate the impact of donor inoculum level on the conjugation frequency, one set of plants were also inoculated with the mixture of donor to recipient to helper of 1:1:1 and 100:1:1. For environmental stress based experiments (section 3.4), the ratio used was 50:1:1 to prevent the inhibition of donor cell growth from the imposed stress by salt and PEG. The inoculated plants were moved to the growth chamber and the bacterial cells were allowed to grow and conjugate for 24 hours under the same conditions the plants were grown previously. The workflow for this experiment is shown in **Fig. 2**.

**Fig. 2.**
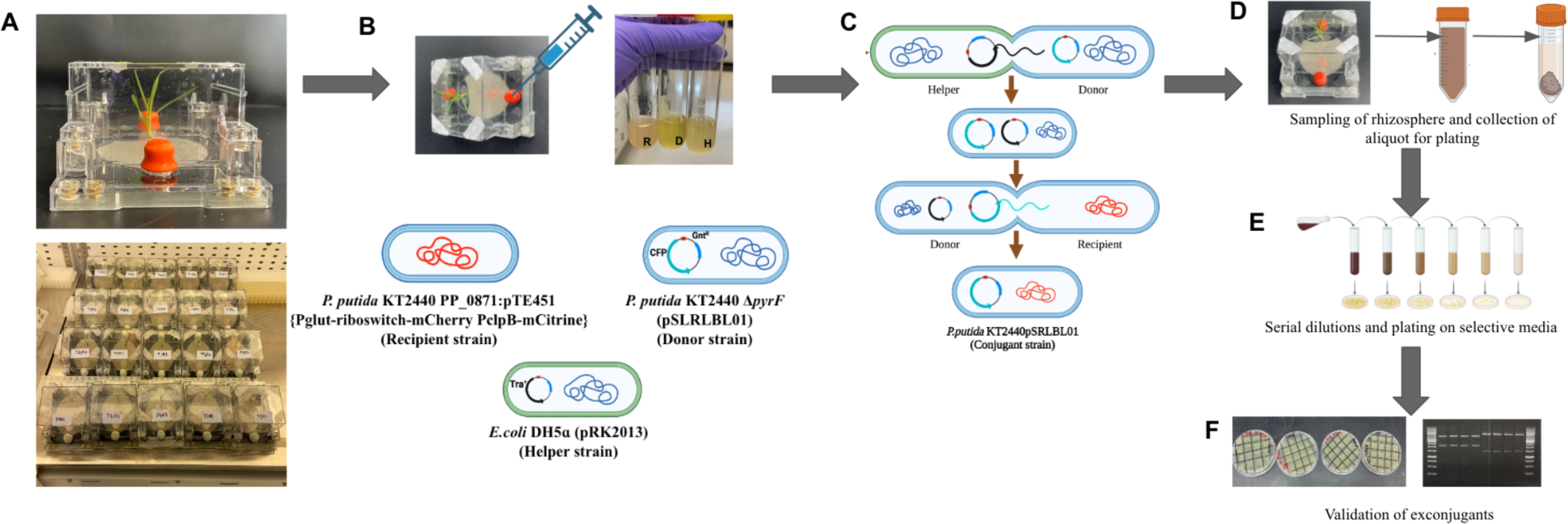
Experimental workflow for quantification of conjugation frequency in the rhizosphere of *B. distachyon* grown in EcoFABs. Growing *B. distachyon* in EcoFABs in plant growth chamber for 2 weeks (A) followed by inoculation of donor (D), recipient (R) and helper (H) strains in the rhizosphere (B). Schematic representation of triparental conjugation system (C). Sampling of rhizosphere 24 hours post-inoculation, dissolved in 5mL PBS, vortexed for 5 min, 1 mL aliquot is collected after sand is settled (D), Serial dilution and plating of 100 µL aliquot on selective media (E), validation of exconjugants by streaking on selective media and restriction enzyme digestion of isolated donor plasmid from exconjugants (F). This figure was created with <u>Biorender.com</u>.

### 2.7 Conjugation experiment on the LB agar plates

To compare the conjugation frequency in nutrient dynamic conditions of the rhizosphere with the optimum nutrient conditions for bacteria, triparental mating was performed on LB agar plates supplemented with 25 µg/mL of uracil. For conjugation mating on plates, donor, recipient and helper cells were mixed in a microcentrifuge tube in the ratio of 10:1:1 with the total volume of 100 uL and was plated on LB with uracil plates. The plates were then incubated at 30 ℃ for 24 hours.

### 2.8 Detection and enumeration of conjugants

After 24 hours of incubation, 1 mL of bacterial suspension from the rhizosphere of the plants in EcoFABs was collected as described above (section 2.2) and serially diluted using PBS. From each dilution, 100 uL was spread on to M9 agar plates, M9 agar plates supplemented with gentamicin and M9 agar plates supplemented with 25 ug/L of uracil to count the number of CFUs/mL of the recipient, donor and conjugants respectively and conjugation frequency was calculated as mentioned below.

The conjugation frequency (CF) was calculated by referring to Equation [17] :

CF = NC/NR

where NC indicates the CFU/mL of the conjugants and NR represents the CFU/mL of recipients.

### 2.9 Graphical representation and statistical analysis of data

GraphPad Prism 9 (GraphPad Software, San Diego, California USA) was used to make all the graphical figures and perform statistical analysis. Phylogenetic trees were constructed using MEGA11 software [27] using maximum likelihood method and were visualized using iTOL [28]. Significant differences were assessed using student’s *t*-test and one-way ANOVA [29]. At a 95% confidence interval, a *p*-value ≥ 0.05 was considered insignificant. In the figures, ** indicates high statistical significance (*p* < 0.01), * indicates statistical significance (*p* < 0.05).

### 2.10 Microscopic imaging of plant roots

The images of the inoculated roots were taken using EVOS M7000 fluorescence microscope (Thermo Fisher Scientific, Waltham, MA) using objectives with 2X and 40X magnification. For imaging purposes, we replaced the plasmid in the donor strain with p101mNeonGreen [30] with constitutively expressed neon green fluorescence. Both donor (green) and recipient (mCherry red) were inoculated in the *B. distachyon* rhizosphere grown for 2 weeks and images were taken 24 hours post-inoculation. For making the images to accommodate color vision deficiency (CVD), we switched the red color of mCherry to magenta using Fiji software [31], the original images are also shown in **Fig. S6**.

## 3. Results and Discussion

### 3.1 Detection and quantification of horizontal gene transfer via conjugation in rhizosphere of *B. distachyon*

Triparental mating under laboratory conditions has been reported for many gram-negative microbes and served to develop a model system for our study. We initiated this project by first demonstrating the desired HGT system with *P. putida* and selection for growth with an antibiotic marker that would enable detectable plasmid transfer events under optimal conditions. A recipient *P. putida* strain could grow in the presence of gentamicin if it acquired a plasmid harboring a *aacII* gene (conferring gentamicin resistance) from a donor strain by a triparental conjugation system. All three strains (donor, recipient and helper) were spotted on a solid agar plate supplemented with LB medium and 25 ug/mL uracil. If the recipient strain successfully acquired the plasmid from the donor strain, we could select for this event by plating the mixture onto M9 glucose plates with gentamicin. The *P. putida* donor strain contained a Δ*pyrF* mutation rendering it unable to grow in minimal media without uracil supplementation, which we use as a counterselection against the donor strain in calculating HGT events. The total number of exconjugants observed were 4.8*10^6^ CFUs per mL with a conjugation frequency of 3*10^−2^ (**Fig. 3A and B**). To further demonstrate HGT in the rhizosphere, we inoculated the three strains in the rhizosphere of *B. distachyon* grown in EcoFABs and plated the conjugants after 24 hours. In the *B. distachyon* rhizosphere, the total number of exconjugants observed were 167 CFUs per mL with a conjugation frequency of 1.4*10^−6^ (Fig. 3A and B).

**Fig 3.**
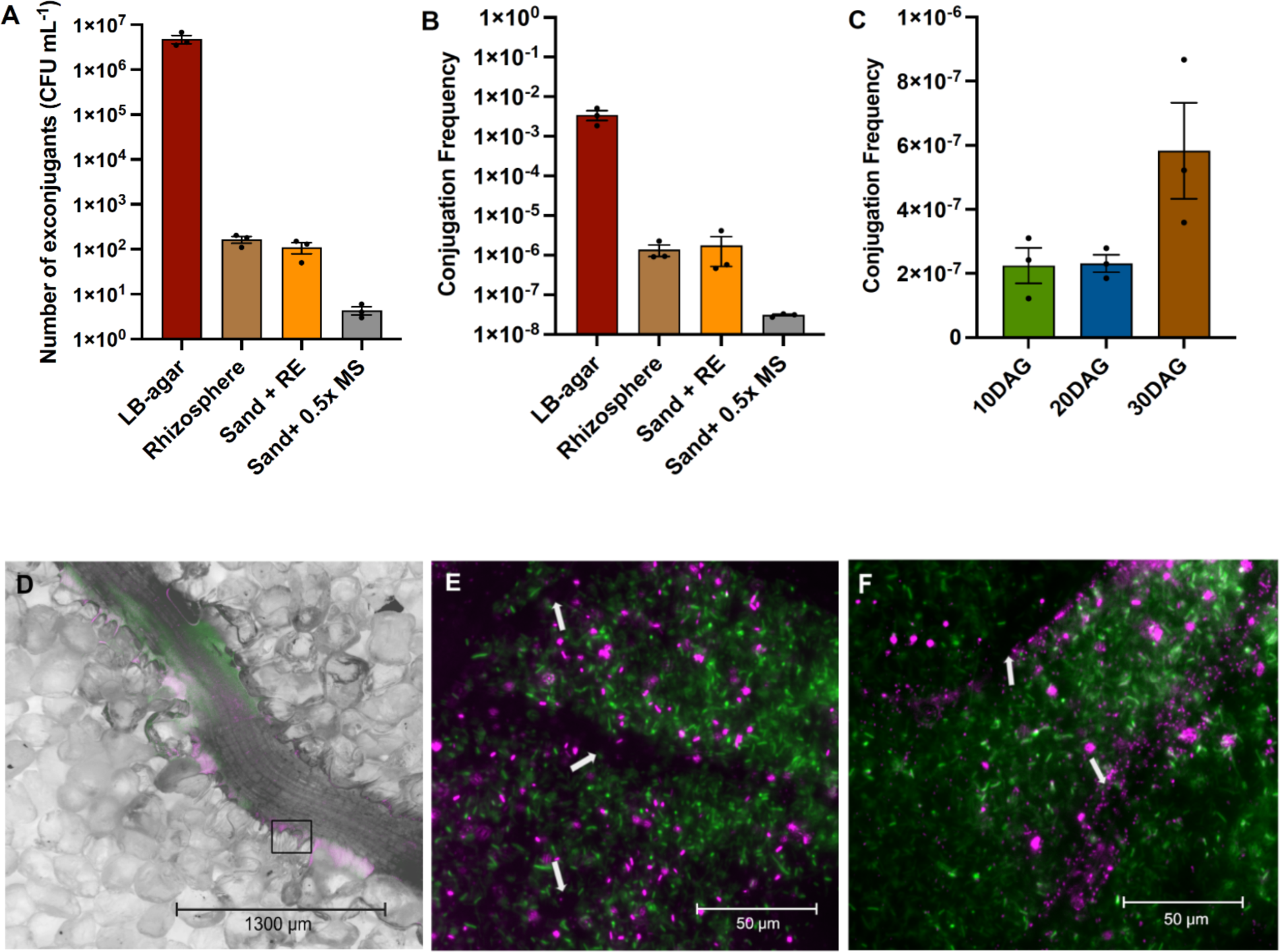
The number of exconjugants (CFU per mL, A) and the conjugation frequency (B) observed with different treatments. RE: root exudate. Conjugation frequency observed at different developmental stages of plant (C) DAG: days after germination. The error bars show the mean standard error of three replicates. Microscopic images of *B. distachyon* roots inoculated with proxy donor (green) and recipient (magenta) strains in EcoFABs at using 2X (D) and 40X (E, F) objectives. The black box in D represents the area magnified for E and F and white arrow in E and F represents the root hair. The red color of mCherry has been replaced by magenta to represent recipient strains.

To our knowledge, this is the first demonstration of HGT using an EcoFAB. The frequency of HGT in the rhizosphere in EcoFABs was lower than that on agar plates, potentially due to fluctuating nutrient conditions of the rhizosphere [32]. The number of conjugants and conjugation frequency in the rhizosphere observed in this study was similar or higher than other studies [33]. Several studies have previously confirmed HGT in soil under various conditions using different soil (without plants) microcosms [34–37]. However, there are very few studies in recent years that have shown HGT in a rhizosphere using a lab-scale microcosm due to the complexity of recapitulating dynamic conditions of a plant associated environment as well as the risks associated with using engineered systems [10,38]. Our results provide evidence demonstrating the EcoFABs as a platform to examine the fate and migration of target plasmids/genes in plant associated environments.

### 3.2 Impact of root exudates and plant developmental stages on frequency of HGT in rhizosphere

We examined the influence of the presence of root exudates on the conjugation frequency in the rhizosphere by comparing the conjugation frequency in the rhizosphere (with plant) to that of EcoFABs filled with sand only (Sand + 0.5 x MS). We also tested if removal of the plant root influences this process when root exudate is still present (Sand + RE) to understand if the presence of root surface is an essential requirement for the HGT process in the rhizosphere. The procedure of removing the plants and adding back the root exudates in the sand filled EcoFABs is described in detail in section 2.2.

The conjugation frequency decreased dramatically in the EcoFABs with only sand (Sand + 0.5x MS) compared to the rhizosphere (**Fig. 3A and B**). This indicated that the compositional change in the rhizosphere due to the presence of plant root exudates could be a major facilitator of HGT in the rhizosphere environment. However, we observed no significant difference in the conjugation frequency between the rhizosphere and the EcoFABs filled with root exudates only (Sand + RE). This clearly indicates that the root exudate is an essential requirement for uptake of plasmids and the presence of the plant root surface may not be an essential factor determining the conjugation frequency. Root exudates provide the necessary nutrients for improved growth and microbial activity which may stimulate the uptake and conjugal transfer of the plasmid [11,33,39]. Additionally, we did not find any major differences in the bacterial density under these two conditions (Rhizosphere and Sand + RE) while the density of donor bacteria was several times lower in the EcoFABs containing only 0.5x MS media **(Fig. S1)**. The microscopy images also suggest that the bacteria (both donor and recipient) are primarily present in the vicinity of roots and root hair where most of the root exudates are released (**Fig. 3D to F**). These results suggest that the root exudate may be an essential factor for determining the conjugation frequency mainly by influencing the bacterial growth and survival in the rhizosphere.

We examined the impact of different plant developmental stages on the conjugation frequencies as it may impact the amount and composition of the root exudates [40]. The results showed that the frequency increased by ∼3X in the plant of later stages (30 DAG) compared to early phases of growth (**Fig. 3C**). This could be due to higher amounts of total root exudates in the plants at later developmental stages leading to higher root colonization by bacteria (donor) and thus stimulating the HGT **(Fig. S2)** [10,11,39]. Molbak et al. (2007) reported that the amount of root exudates and root growth are the key parameters impacting plasmid transfer in the rhizosphere while the root surface features did not play an important role in this process. They compared the rates of horizontal plasmid transfer in the rhizosphere of pea and barley and showed that pea roots had 10 times higher transfer rate due to higher rate of root exudation of pea roots which increased the colonization of donor strains [10]. To understand the changes in the composition of carbon substrates in root exudates, we quantified the composition of organic acids and sugars possibly present in the root exudates of *Brachypodium* at all the three stages of the plant [41]. The total amount of organic acids detected (malic acid, citric acid and succinic acid) were higher than that of sugars (glucose and xylose) at all stages of plant growth up to 30 DAG **(Fig. S3)**. Several studies have confirmed that organic acids in the plant root exudates play a major role in inducing chemotaxis and therefore are crucial for root colonization by bacteria in the rhizosphere [42–44]. In our study, the concentration of one specific organic acid (malic acid) increased proportionally with the plant developmental stages and could be correlated with the increased conjugation frequency at 30 DAG **(Fig. S3)**. This was observed in a study that certain organic acids from plant root exudates stimulated the plasmid (pFG4) transformation in *Acinetobacter sp.* BD413 in soil [45]. However, additional direct testing of these metabolites is required to completely understand the impacts of different root exudate components on conjugation efficiency in the rhizosphere.

### 3.3 Measuring impact of donor to recipient ratio on conjugation frequency in rhizosphere

Conjugation requires physical contact between donor and recipient, therefore the ratio of donor to recipient (RD/R) cells has been identified as one of the most critical factors that determine the frequency of conjugal transfer in several studies [17,46]. To understand the impact of inoculum size of the donor strain on conjugation frequency in the rhizosphere, we used 3 different ratios of donor to recipient 1:1, 10:1 and 100:1. Results showed that increasing the inoculum size of the donor by 10X or 100X significantly improved the conjugation frequency in the rhizosphere by ∼30X and ∼200X respectively (**Fig. 4**). Lampkowska et al. (2001) published a standardized and optimized protocol for conjugation between two lactococcal strains and found that the frequency increased with increasing the ratio from 1:10 to 1:1 [46]. However, in a recent study Shi et al. (2023) reported that the smaller the RD/R, the higher the conjugant concentration [17]. It is therefore possible that the impact of RD/R is variable under different experimental conditions and may also depend on the colonization rate or fitness of the two bacterial strains in the rhizosphere environment. In the case examined in the present study, the donor was generally found to be lower in numbers than the recipient strain under all conditions **(Fig. S1 and S2 and S4)** possibly due to its dependency on the presence of uracil for survival and therefore increasing the ratio of RD/R improved the conjugation efficiency.

**Fig 4.**
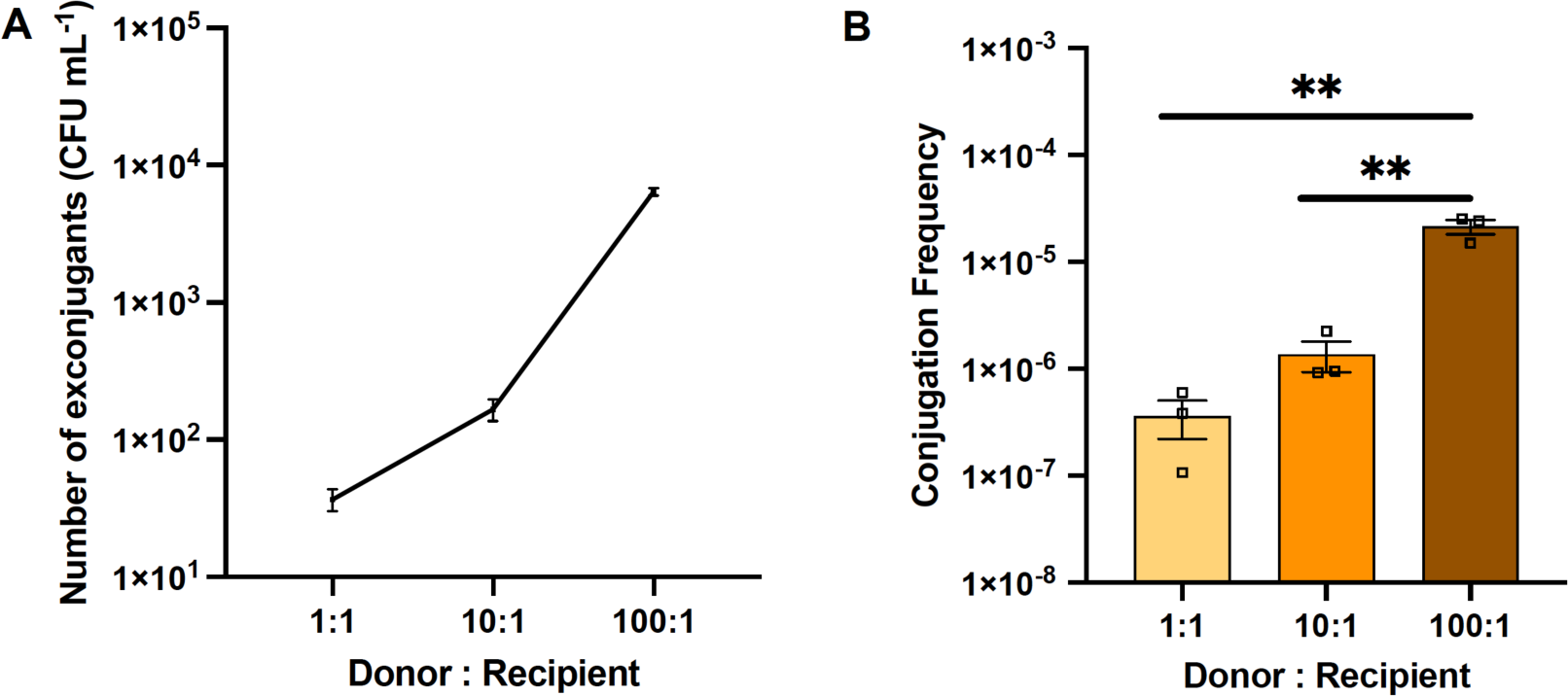
Change in the number of exconjugants (CFU per mL, A) and the conjugation frequency (B) with increasing the donor to recipient ratio. The error bars show the mean standard error of the three replicates. The two asterisk (**) indicates p value < 0.005 when the student’s t-test was performed.

### 3.4 Influence of environmental stress on the conjugation frequency in the rhizosphere

Although there are few studies that have tested the impact of rhizosphere environmental conditions on HGT frequencies, two studies have shown that physical conditions of soil such as soil pH, moisture level, soil type or the mineral composition have an influence on the transfer of plasmids [16,17]. Drought and other associated changes in the rhizosphere could be another important parameter that influences HGT frequency, however it has not yet been investigated in detail. To evaluate the impact of several environmental stress conditions on HGT in the rhizosphere, we examined the effects of osmotic stress by exogenously adding salt and PEG in the 0.5x MS medium (separately) at two different concentrations into the EcoFABs during the seedling transfer stage (see section 2.1). Results indicate that the number of exconjugants was highest in the rhizosphere with 10% PEG with ∼1.5*10^4^ CFUs/mL while higher concentrations of PEG (20%) decreased the conjugation frequency compared to the no-stress controls. Addition of salt to the rhizosphere decreased the number of conjugants fourfold with both moderate and high levels of salt content as compared to the rhizosphere with no stress compounds (**Fig. 4**).

Our results show that the conjugation frequency may vary with the environmental conditions in the rhizosphere; it can be increased if the plant is exposed to a moderate (sub-inhibitory) level of drought/osmotic stress and conversely decreases in the presence of salt even at non-inhibitory concentrations. It was previously reported that high levels of stress reduce the conjugation frequency due to the inhibition of cell activities. Contrary to this, sub-inhibitory levels of stress might activate the global stress response mechanisms in bacterial cells e.g., biofilm formation or increased membrane permeability creating favorable conditions for conjugation in the rhizosphere [47,48]. This is consistent with the report by de la Rosa et al. (2021), sub-inhibitory levels of antibiotics were found to increase the conjugation frequency due to activation of SOS response genes [49]. Addition of moderate and high concentrations of salt to the rhizosphere substantially decreased the number of exconjugants by ∼4X as compared to the rhizosphere containing no stress compounds (**Fig. 5**). This could be mainly due to the inhibitory effect of salt on donor cells **(Fig. S4)**. This is supported by Tan et al. (2019) who reported that an increase in salinity in soils hindered the conjugation frequency of plasmids containing antibiotic resistance genes (ARGs), mainly due to decreased fitness of donor cells under such conditions [50].

**Fig 5.**
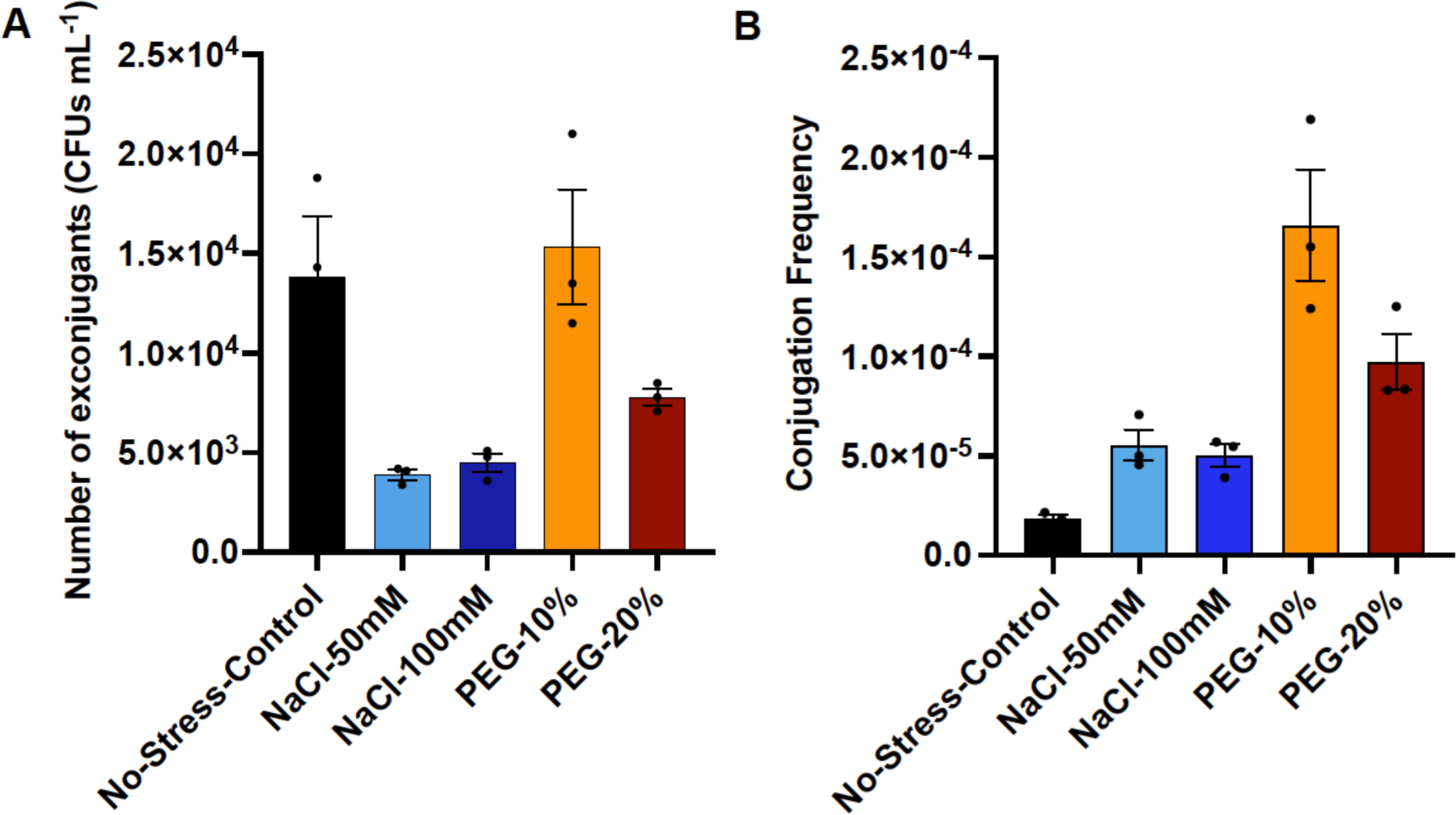
The number of exconjugants (CFU per mL, A) and the conjugation frequency (B) observed under salinity and osmotic stress conditions. The error bars show the mean standard error of the three replicates.

Thus, several factors in the soil or rhizosphere may influence the efficiency with which microbes take up plasmids. As such, it is important to consider multiple parameters while investigating the fate of plasmid or genes intentionally or unintentionally disseminated into a field and or plant-associated environment.

### 3.5 Detection of HGT between bacteria of different genera in rhizosphere

To determine whether the plasmid of one bacterial species in the rhizosphere is also taken up by bacteria of other species or genera, we performed conjugation experiments in the EcoFABs with a plant-associated bacterium *Burkholderia sp.* OAS925 as the recipient. *B. sp.* OAS925 has been shown previously to be an efficient colonizer of the *Brachypodium* rhizosphere [9].

Our results indicate that *Burkholderia* can also take up the plasmid from other bacterial genera efficiently and the frequency of conjugation increases under stressed environmental conditions such as salinity. We observe that under normal (no-stress) conditions, the number of exconjugants in the rhizosphere were ∼ 600 CFUs per mL when *B. sp.* OAS925 was used as recipient and *P. putida* as the donor of the plasmid and was 100X lower than observed with the recipient of the same species (**Fig. 5A and 6A**). Addition of salt in the medium increased the intergeneric transfer of plasmid and the highest frequency of ∼10^−2^ was observed in the rhizosphere supplemented with 100mM Salt (**Fig. 6B**). The cell densities of two bacteria in the salt added media were lower than no-stress conditions under both high and low NaCl levels indicating a negative impact of salt on the growth of both recipient and donor. This shows that the stress due to high salt content could trigger the plasmid transfer between different bacterial genera in the rhizosphere.

**Fig 6.**
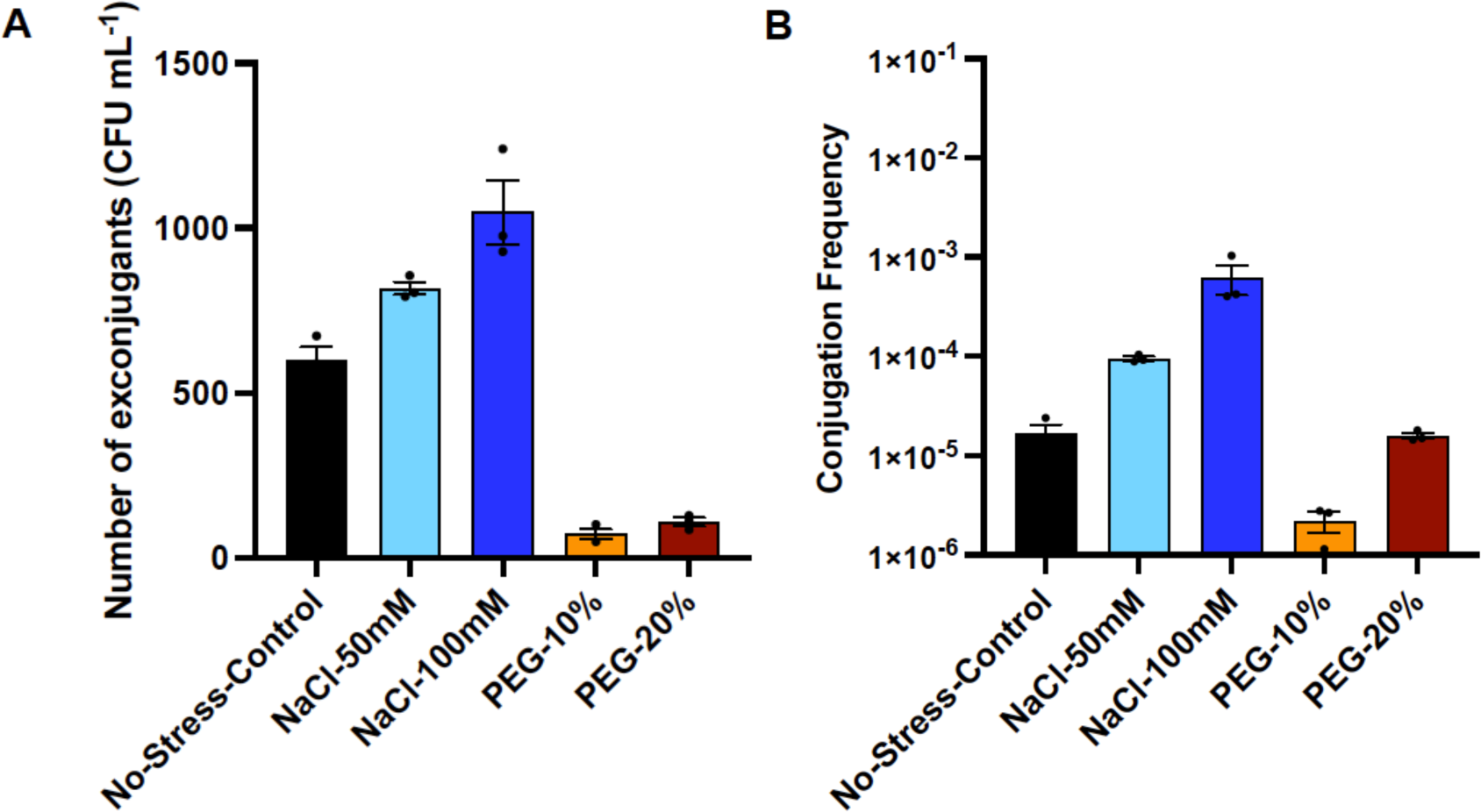
The number of exconjugants (CFU per mL, A) and the conjugation frequency (B) observed when *Burkholderia sp.* OAS925 was used as a recipient strain. The error bars show the mean standard error of the three replicates.

Osmotic changes can also be exogenously induced using PEG and was implemented using 10% and 20% PEG as described in methods. Increase in osmotic levels by adding PEG in the medium lowered the number of exconjugants and conjugation frequency. However, while the density of donor and recipient decreased at the higher concentration of PEG (20%), it was comparable to no stress conditions under lower PEG (10%) concentration **(Fig. S5)**. This suggests that the final conjugal transfer frequency under these conditions is potentially a trade-off between the number of cells and the stress that may induce gene transfer. Since PEG causes both cell growth inhibition and stress there may be an intermediate level between 10% and 20% where conjugal transfer frequency is similar or higher to no stress conditions.

These results provide evidence of possible transmission of plasmids between two different bacterial genera in the rhizosphere microbial community under both normal and abiotic stress conditions. It has been shown previously that bacteria of different genera can conjugate and transfer the plasmid in order to survive in stress conditions [18,51], however, it has not extensively studied in the rhizosphere. Further studies will reveal the fate of plasmids in the presence of a complex rhizosphere microbial community mimicking the natural conditions of the rhizosphere and what factors may influence the frequency of HGT.

### 3.6 Influence of self-transmissible plasmids in the rhizosphere on the conjugation frequency

The conjugation process in a natural environment usually involves a donor carrying a self-transmissible conjugative plasmid that is transferred to other bacteria (recipients) in the vicinity. These plasmids have a transfer (*tra*) genes cluster which encodes all the proteins required for their efficient transfer from donor to recipient [33]. In some cases including this study, these self-transmissible plasmids also facilitate the transfer of other non-conjugative mobilizable plasmids (triparental conjugation) [52]. The function of all the *tra* genes and the transfer mechanism has been explained in detail in several studies [52–55]. Very few bacterial species in the rhizosphere contain the conjugative plasmids with *tra* genes cluster as shown in the phylogenetic tree of commonly observed bacteria in the rhizosphere of *B. distachyon* and other related monocot grasses (**Fig. 7A**). Therefore, the frequency of gene transfer within rhizosphere bacterial communities may be at low levels and may vary on the number of bacteria carrying these self-transmissible plasmids. While it is plausible that bacterial strains with *tra* genes or genes encoding similar functions could facilitate a triparental gene transfer in a natural environment, our review of literature did not reveal any such examples. Exchange of genetic material between two strains is more commonly documented, and the majority of the studies on horizontal gene transfer via conjugation involves self-transmissible conjugative plasmids [10,33,38]. The donor plasmid (pSLRLBL01) used in this study is a mobilizable plasmid and does not have the *tra* gene cluster. To understand the possibility of plasmid transfer without the help of *tra* genes, we performed biparental mating on LB medium excluding the helper strain.

**Fig 7.**
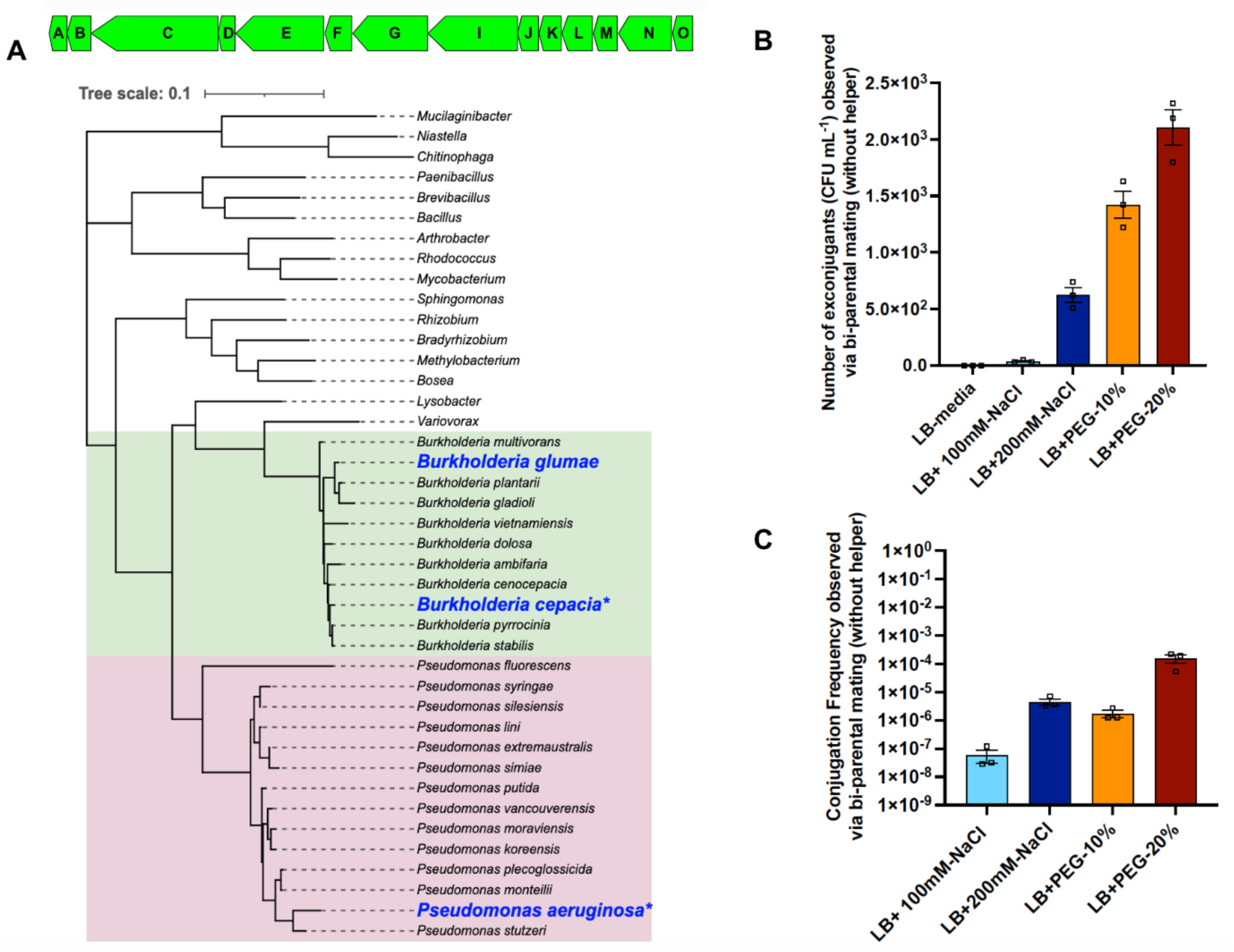
Phylogenetic tree of dominant bacteria in rhizosphere of *Brachypodium* and related monocot plants [56–58], the blue colored bacteria contain *tra* genes cluster (shown on the top of the tree) present in the helper plasmid pRK2013, the letters A, B, C, D, G, I, J, K, L, M, N and O represents genes *traA, traB, traC, traD, traE, traF, traG, traI, traJ, traK, traL, traM, traN* and *traO* respectively, * represents that the *tra* gene cluster is on a plasmid, the highlighted regions are the genera for which specific species found in the rhizosphere are shown (A). The number of exconjugants (CFU per mL, B) and the conjugation frequency (C) observed via biparental mating on LB medium under salinity and osmotic stress conditions. The error bars show the mean standard error of the three replicates.

No exconjugants were observed on LB medium plates under optimum conditions. However, addition of salt or PEG in the media led to detectable plasmid transfer and the highest frequency of ∼10^−4^ was observed in the LB medium with 20% PEG. This frequency is several orders of magnitude lower than we observe in the triparental mating condition on LB medium agar plates (**Fig. 3B**). No exconjugants were observed when the experiment was conducted in the rhizosphere environment, likely due to the overall decrease in efficiency between HGT compared to the agar plate setup and the soil environment. Additional experiments are required to further examine if transfer of mobilizable plasmids is possible in the absence of a conjugative helper plasmid, and if this can be observed in a plant root associated environment. Environmental stressors alone may enhance HGT rates to detectable levels without a strict requirement for a helper strain. These observations indicate that it may be useful to examine HGT frequencies in microbial communities that may not have self-transmissible plasmids.

### 3.7 Conclusions

EcoFABs provide an efficient and useful platform for assessing HGT frequency in the rhizosphere as they can simulate highly variable conditions of a rhizosphere environment. We observed the plasmid transfer under both normal and environmental stressed conditions and investigated different parameters that may influence this process. The ratio of donor to recipient had a major impact on the conjugation frequency and may vary due to the difference in the fitness of the donor and recipient bacteria. The developmental stage of the plant and the root exudate composition also influence this frequency. We have also shown the possibility of conjugation between *P. putida* and *Burkholderia* sp. providing evidence that some of the commonly found bacteria in the rhizosphere can efficiently take up plasmids from other microbial genera. In addition, we observed that intraspecies plasmid transfer is increased in the conditions of moderate osmotic stress while saline conditions facilitate the conjugation between bacteria of different genera. This study underscores the multi-parametric nature of the rhizosphere with dynamic enhancing and inhibiting conditions that can impact HGT. Therefore, a controlled way to modulate multiple factors to thoroughly investigate the gene transfer process is required for which EcoFABs could be a very useful tool. Presence of self-transmissible plasmids may not be necessary for the HGT process under stressed conditions; however, this is yet to be investigated rigorously in a rhizosphere environment. This study provides sufficient evidence to confirm the HGT process under various conditions in the rhizosphere that can be quantified using EcoFABs and therefore precautionary measures are needed when disseminating engineered microbes containing mobilizable plasmids in the field or in agricultural environments.

## Acknowledgements

The authors thank the Mukhopadhyay group and the Microbial Community Analysis & Functional Evaluation in Soils (m-CAFEs) team for their constructive feedback regarding this manuscript. We thank John Vogel for providing *Brachypodium distachyon* BD21-3 seeds and Lorenzo Washington for providing the plasmid (p101mNeonGreen) for imaging purposes. We also thank Jonathan Diab for their help with initial training with EcoFAB and osmotic stress assays and Venkataramana Pidatala for their help with HPLC data acquisition and analysis. This project was supported by m-CAFEs, a Science Focus Area led by Lawrence Berkeley National Laboratory based upon work supported by the US Department of Energy, Office of Science, Office of Biological & Environmental Research under contract number DE-AC02-05CH11231.

## Author Contributions

SP and AM conceptualized the study. SP and SR designed and conducted the experiments. PFA built and provided the EcoFABs to perform the experiments. Microscopy and Imaging were performed by SP, HHL and JCM. TE and AM provided the supervision. AM, JCM and TRN acquired the funds for the project. SP drafted the initial manuscript. All authors have read, provided feedback, and approved the manuscript for publication.

