## Supplementary Information for "Assessing horizontal gene transfer in the rhizosphere of *Brachypodium distachyon* using fabricated ecosystems (EcoFABs)"

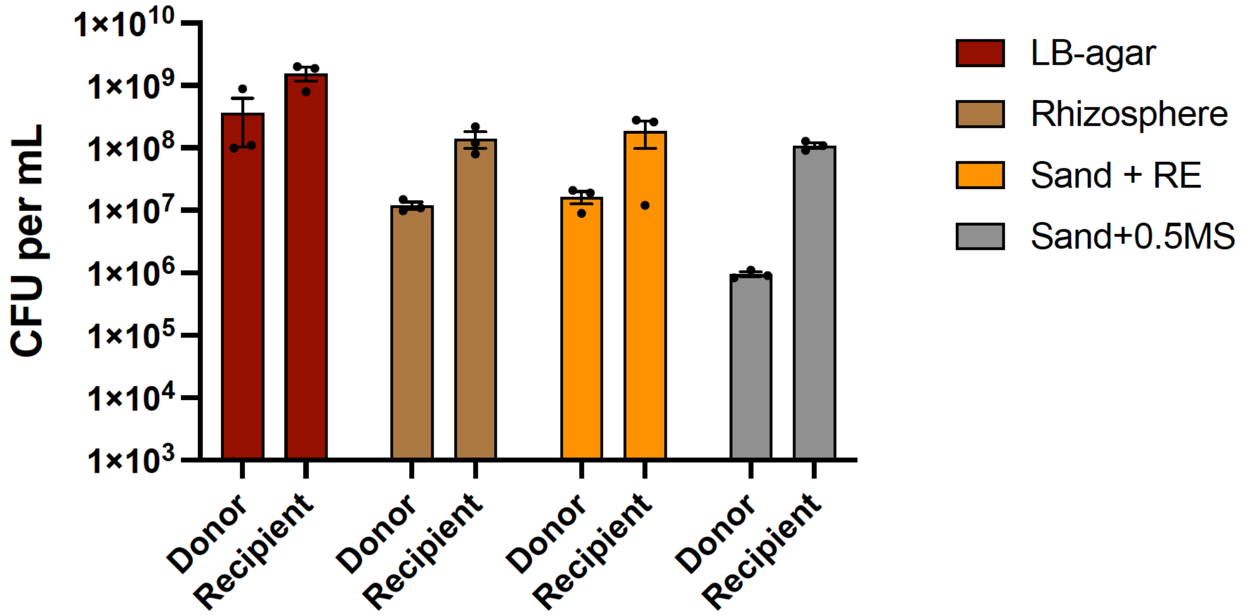


Fig S1. The number of donor and recipient (CFU per mL) observed under different treatments. The error bars show the mean standard error of the three replicates.

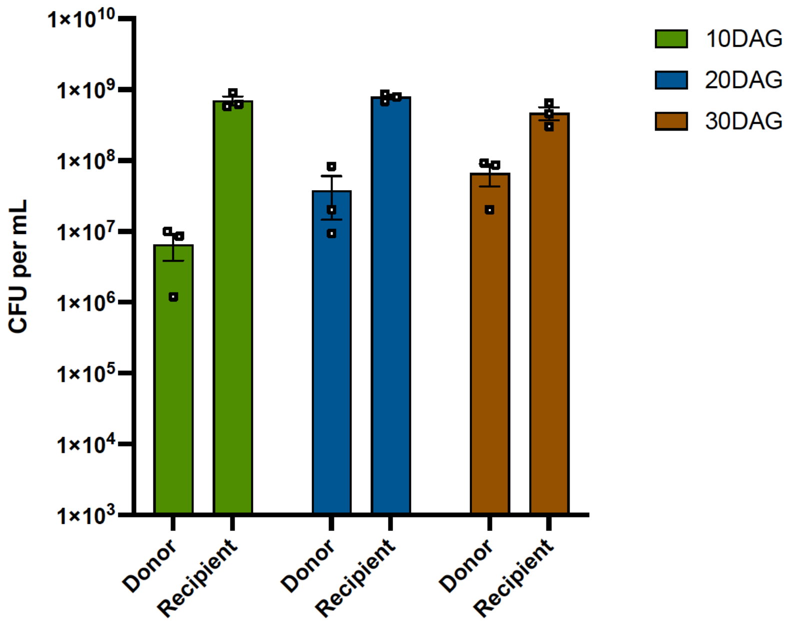


Fig S2. The number of donor and recipient (CFU per mL) observed at different developmental stages of the plant. DAG: days after germination. The error bars show the mean standard error of the three replicates.

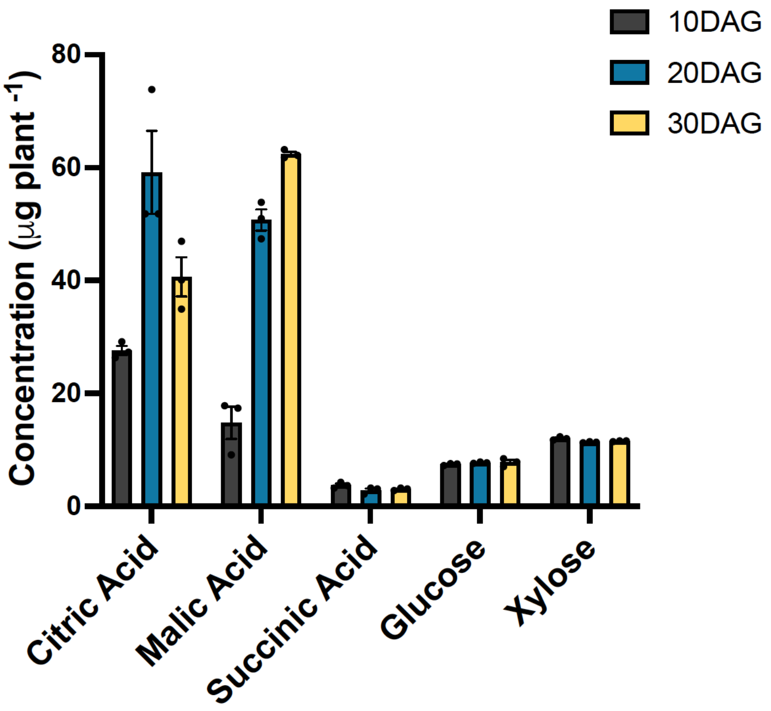


Fig S3. Concentration of organic acids and sugars detected in the root exudates extracted from BD21-3 at 10DAG, 20DAG and 30DAG. DAG: days after germination. The error bars show the mean standard error of the three replicates.

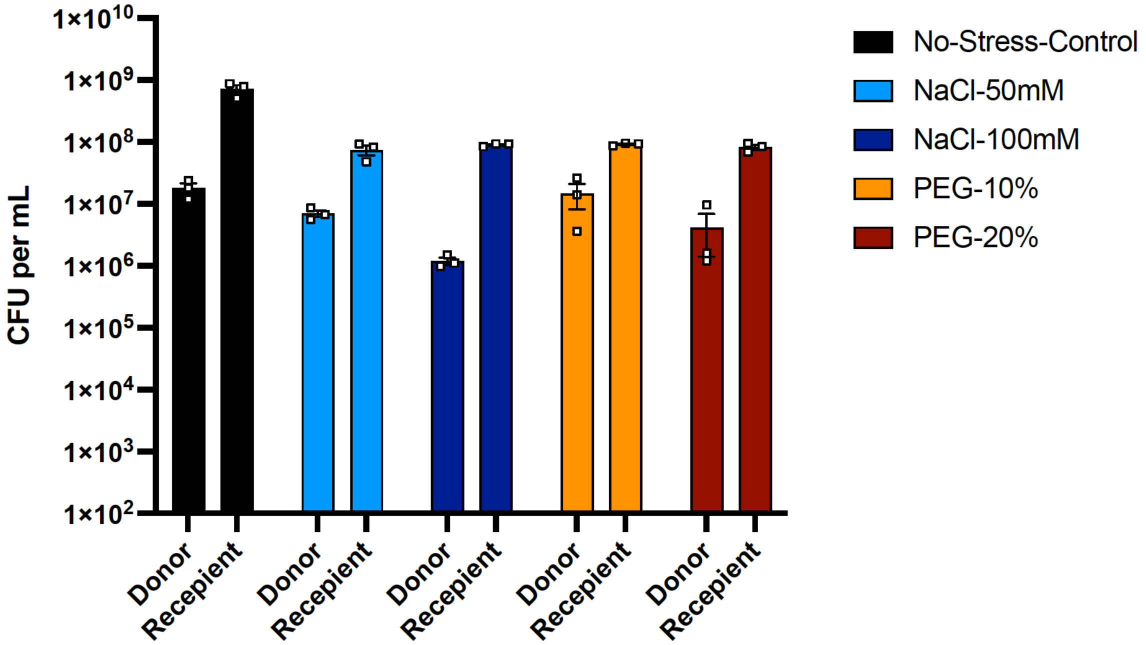


Fig S4. The number of donor and recipient (CFU per mL) observed under different treatments. The error bars show the mean standard error of the three replicates.

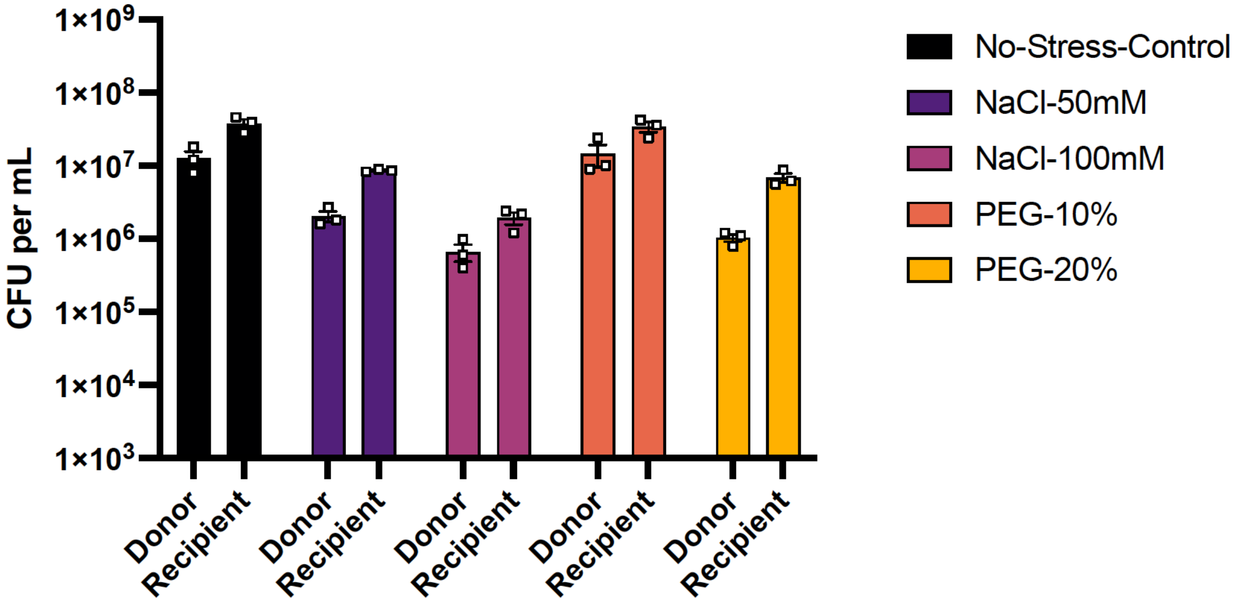


Fig S5. The number of donor (*P.putida* KT2440Δ*pyrF)* and recipient (*B. sp.* OAS925)(CFU per mL) observed under different treatments with intergeneric conjugation in the rhizosphere. The error bars show the mean standard error of the three replicates.

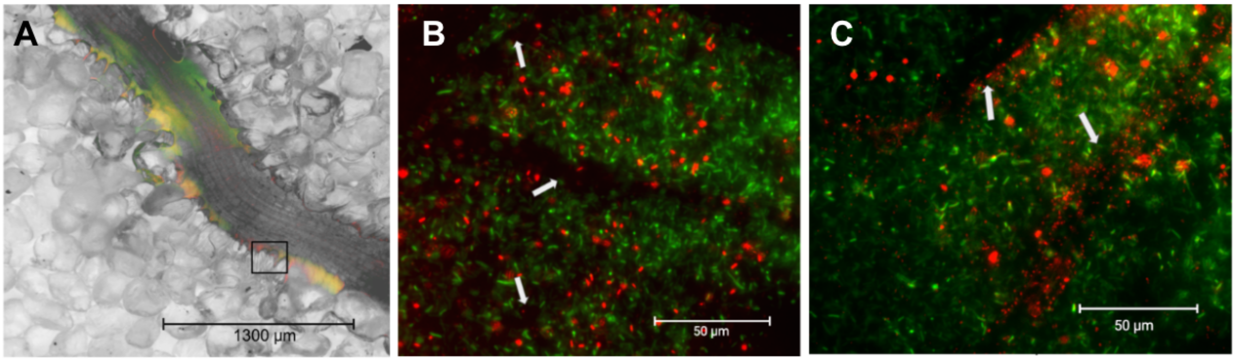


Fig S6. Microscopic images of *B. distachyon* roots inoculated with proxy donor (green) and recipient (red) strains in EcoFABs using 2X (A) and 40X (B, C) objectives. The black box in image A represents the area magnified for images B and C. White arrows in B and C represent the root hair. This is the original version of the image presented in Fig 3D-3F where red of mCherry is replaced by magenta color.
